# MATRIEX Imaging: Multi-Area Two-photon Real-time In-vivo Explorer

**DOI:** 10.1101/510545

**Authors:** Mengke Yang, Zhenqiao Zhou, Jianxiong Zhang, Tong Li, Jiangheng Guan, Xiang Liao, Bing Leng, Jing Lyu, Junan Yan, Kuan Zhang, Yan Gong, Yuguo Tang, Zhiming Zhu, Zsuzsanna Varga, Arthur Konnerth, Jinsong Gao, Xiaowei Chen, Hongbo Jia

## Abstract

Two-photon laser scanning microscopy, originally developed since 1990s^1^, has been widely applied for biomedical research in recent decades, particularly popular among neuroscientists for studying neural functions *in vivo*^2^. However, it is typically restricted to one imaging area that is orthogonal to the optical axis. Here, we demonstrate a novel multi-axis optical conjugation method that enables two-photon imaging at single-cell resolution simultaneously in multiple areas at different depths, each of which could have a view diameter of ~200 μm and could be largely freely targeted within a zone up to 12-mm diameter. For example, we show simultaneous imaging of neuronal activities in the primary visual cortex (V1), the primary motor cortex (M1) and the hippocampal CA1 region of awake mice. This method can be readily implemented on a single conventional two-photon microscope to enable multi-area exploration of neuronal activities *in vivo*.

---

A unique advantage of two-photon (and three-photon^3^) versus single-photon fluorescence imaging technologies for studying neural activities in intact living brains is its accuracy in highly scattering and densely labeled brain tissues^4^, acquiring the images pixel-by-pixel-wise with minimal crosstalk and thus without the necessity of deconvolution or other complex computational reconstruction *post-hoc*. Unfortunately, this major advantage induces also a major drawback, i.e., the area of tissues that can be simultaneously imaged while maintaining single-cell resolution is highly restricted. Many methods have been successfully established to extend such limit. For example: on the axial dimension, using electrically tunable lens to remotely drive the focal depth^5, 6^, or using Bessel-shaped laser beam with prolonged focal point-spread-function^7^; on the lateral dimension, using large-caliber microscope optics^8^ with multiplexed subfield scanning techniques^9–12^, or combining two or more scanning rigs together onto one animal^13^, or using fast rotating devices for alternating the imaging field-of-view^14^. However, most of these methods require highly complex mechanic and electronic devices as well as highly customized software, which are difficult to be implemented for many neuroscience research labs. Here, we choose a different strategy by dealing with not the laser scanning but the optical conjugation. This method, termed as MATRIEX (Multi-Area Two-Photon Real-time In-vivo Explorer), allows the experimenter to target multiple brain areas within a large zone, to achieve imaging at single-cell resolution in each of the areas simultaneously, by means of acquiring images through the original raster scanning and image acquisition software of the conventional two-photon microscope.

As a convention of optical design, most of optical components of imaging systems are arranged along one optical axis. Here, we demonstrate that multi-axis optical conjugation is also feasible in two-photon laser scanning microscopy. Fig. 1a shows the basic concept of the method, whereas the conventional microscope objective of water-immersion and medium or high magnification (typically, 16X - 40X) is replaced by a customized compound objective assembly (Fig. 1a, middle magnified view), consisting of multiple mini-objectives (MOs) that have a lateral magnification factor of ~8X (custom designed, see Suppl. Fig. 1 for details) and a low magnification air objective (AO, 2X - 5X, a broad range of low-cost standard industrial products available). Multiple MOs with different working distances could conjugate multiple imaging areas at different depths onto one virtual image plane under the AO. Thus, a simple raster scanning device could scan through all the imaging areas simultaneously within one frame of image scan, resulting in a single rectangular image with multiple circular subfields.

**Figure 1.**
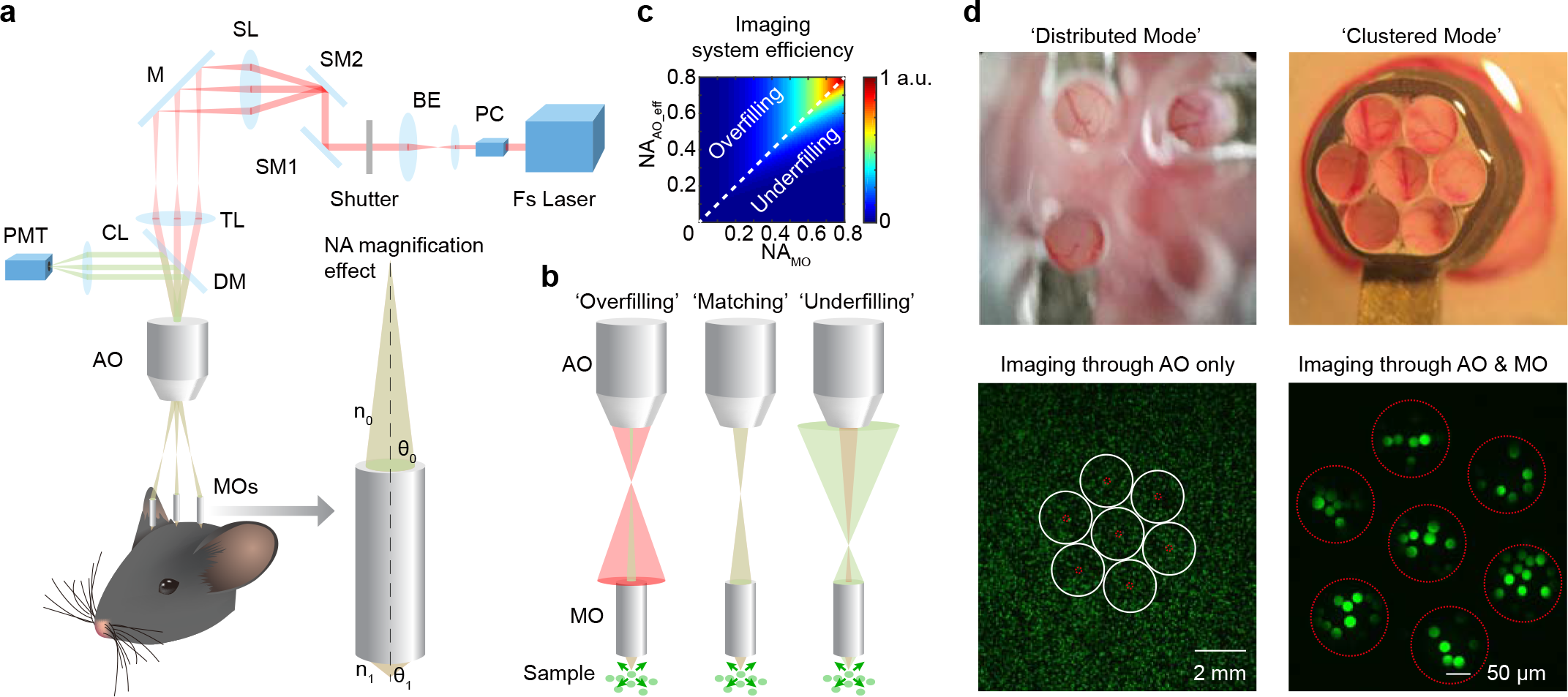
Design and implementation of MATRIEX imaging. **a**, Cartoon diagram of the imaging system and NA magnification effect. The upper stationary part of the imaging system is identical as conventional two-photon microscope system: Fs Laser: Femtosecond Pulsing Laser. PC: Pockels Cell, BE: Beam Expander, SM1 & SM2: Scanning Mirrors 1 & 2 (12-kHz resonant mirror & galvo mirror), SL: Scan Lens, M: Mirror to feed scanning path into microscope tube, TL: Tube Lens, DM: Dichroic Mirror to split excitation and emission light, CL: Collection Lens for fluorescence photons, PMT: Photomultiplier Tube (GaAsP Detector). The lower customized adjustable part of the imaging system contains one AO (Air Objective) and multiple MOs (Miniaturized Objectives). Enlarged inset cartoon show the NA magnification effect, where *θ*_0_, *θ*_1_ are the angles of excitation light cone on both sides of the MO,*n*_0_ = 1, *n*_1_ = 1.33 are the refractive index of air and water, respectively. **b**, Cartoon illustrating three different scenarios of NA conjugation between the AO and the MO. Red shade: excitation light path, Green shade: emission light path. Brown-green shade: the overlay of excitation light and emission light paths. Small green arrows: mimic of the scattering of fluorescence photons. **c**, Total fluorescence throughput calculated based on realistic optical parameters and different combinations of NA values of AO and MO. **d**, Upper two images: pictures taken by smartphone directly towards the animal skull with MOs configured in different modes. Lower two images: demonstration of the two-stage magnification effect by using 20 μm fluorescent beads. White circles on lower-left image indicate the equivalent position of the outer rim of each MO, and red circles show the actual field-of-view underneath each MO.

The trick to achieve a fair resolution here is the effect of numerical aperture (NA) magnification: the relatively low NA value of the excitation light cone after the low-mag AO (around 0.1) at the virtual image plane is magnified by the MO by its angular magnification factor (Fig. 1a, right-side magnified view). Thus, the theoretical effective NA value of the excitation light cone towards the specimen *NA*_*AO*_eff_ is given by the following equation:

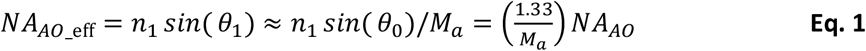

Where *n*_1_ = 1.33 is the refractive index of water, *NA*_*AO*_ is the nominal NA value of the AO, *M*_*a*_ = −*θ*_0_⁄ *θ*_1_= 0.1675 is the angular magnification factor of MO (calculated by the parameters given by the MO manufacturer, see Suppl. Fig. 1 for details). For the different AO models that we have tested, the calculated *NA*_*AO*_eff_ is shown in Table 1.

**Table 1.** NA value of various air objectives tested for the MATRIEX imaging method.

| Air Objective Model | $NA_{AO}$ | $NA_{AO\_eff}$ |
| --- | --- | --- |
| Mitutoyo 2X | 0.055 | 0.44 |
| Olympus 2.5X | 0.08 | 0.64 |
| Olympus 4X | 0.1 | 0.79 |
| Olympus 5X | 0.1 | 0.79 |

However, this theoretical effective NA value for the excitation light cone is limited by the nominal NA value of the MO (*NA*_*MO*_), which is 0.51 in our case. Thus we bring forward three different scenarios (illustrated by Fig. 1b): ‘overfilling’ (*NA*_*AO*_eff__ > *NA_MO_*), ‘matching’ (*NA*_*AO*_eff__ = *NA_MO_*), ‘underfilling’ (*NA*_*AO*_eff__ < *NA_MO_*). Despite a higher NA value is desirable for achieving a higher resolution (smaller focal volume for the fluorescence excitation), mismatching of NA could greatly affect the coupling efficiency at the interface. On the other hand, due to the scattering of fluorescence photons through brain tissue before being collected by the optical assembly, the exact ‘matching’ of NA may not be the ideal scenario. Thus, we calculated look-up heatmaps (Fig. 1c) to show total imaging efficiency upon different combinations of NA values. The ideal NA combination relation to achieve best efficiency deviates from a diagonal line, indicating that a slight ‘overfilling’ is practically more desirable than ‘underfilling’.

There are two general modes for the positioning of MOs: (1) ‘distributed mode’, to place a few (e.g., 3 pieces) MOs that have some spacing between each other and can be individually micromanipulated above several small imaging windows (e.g., Fig. 1d upper-left image); (2) ‘clustered mode’, to bundle many MOs (e.g., 6 pieces around 1 core piece, looks like a revolver gun, Fig. 1d upper-right image) tightly together over one big imaging window. To demonstrate the two-stage magnification effect, we used fluorescent beads (20 μm diameter) to compare the image acquired through only the AO (Fig. 1d lower-left image) and the image acquired through the combined MO+AO in the ‘clustered mode’ (Fig. 1d lower-right image). In practice, for imaging neurons in mammalian brains *in vivo*, the ‘distributed mode’ is more favorable due to flexibility to handle in 3D space, less damage to brain tissue caused by craniotomy and minimal inter-area crosstalk owing to the gap between adjacent MOs. In either mode, the maximum size of targetable zone is limited by the maximum size of scanning field under the AO in the conventional microscope, typically, for an AO of 2X magnification, the diameter of targetable zone is ~12 mm (See also Suppl. Methods).

To demonstrate the application of the MATRIEX method, we performed two-photon Ca^2+^ imaging of GCaMP6f-labeled neurons simultaneously in primary visual cortex (V1), primary motor cortex (M1) and the hippocampal CA1 region (Fig. 2, Supplementary Video 1). The cartoon picture (Fig. 2a) shows the configuration of the three MOs, where the two MOs for V1 and M1 were directly placed above the cortex, and the MO for the hippocampal CA1 region was inserted after surgical removal of the cortical tissue above. The surgical insertion procedure was similar as in previous literature^15, 16^. Note that the MOs were customized with different parameters for the cortex and for the hippocampus, thus they could all conjugate to one virtual image plane under the AO (see Suppl. Fig. 1 for details). On a conventional two-photon microscope equipped with 12 kHz resonant scanner, we scanned the full field of view at 1200 × 1200 pixels and 10 Hz. In the full scanning field, the three areas conjugated by the three MOs were readily visible (Fig. 2b), and we digitally zoomed in each of them to show single cells (Fig. 2c). The spontaneous Ca^2+^ signal traces of all identified neurons were shown in Fig. 2d (220 neurons, total recording duration 300 s, example traces with magnified view shown on Fig. 2e). Neuropil contamination to the neuronal Ca^2+^ signals were nearly absent regardless of their relative position (Suppl. Fig. 2), indicating that the multi-axis conjugation did not affect single-cell resolution.

**Figure 2.**
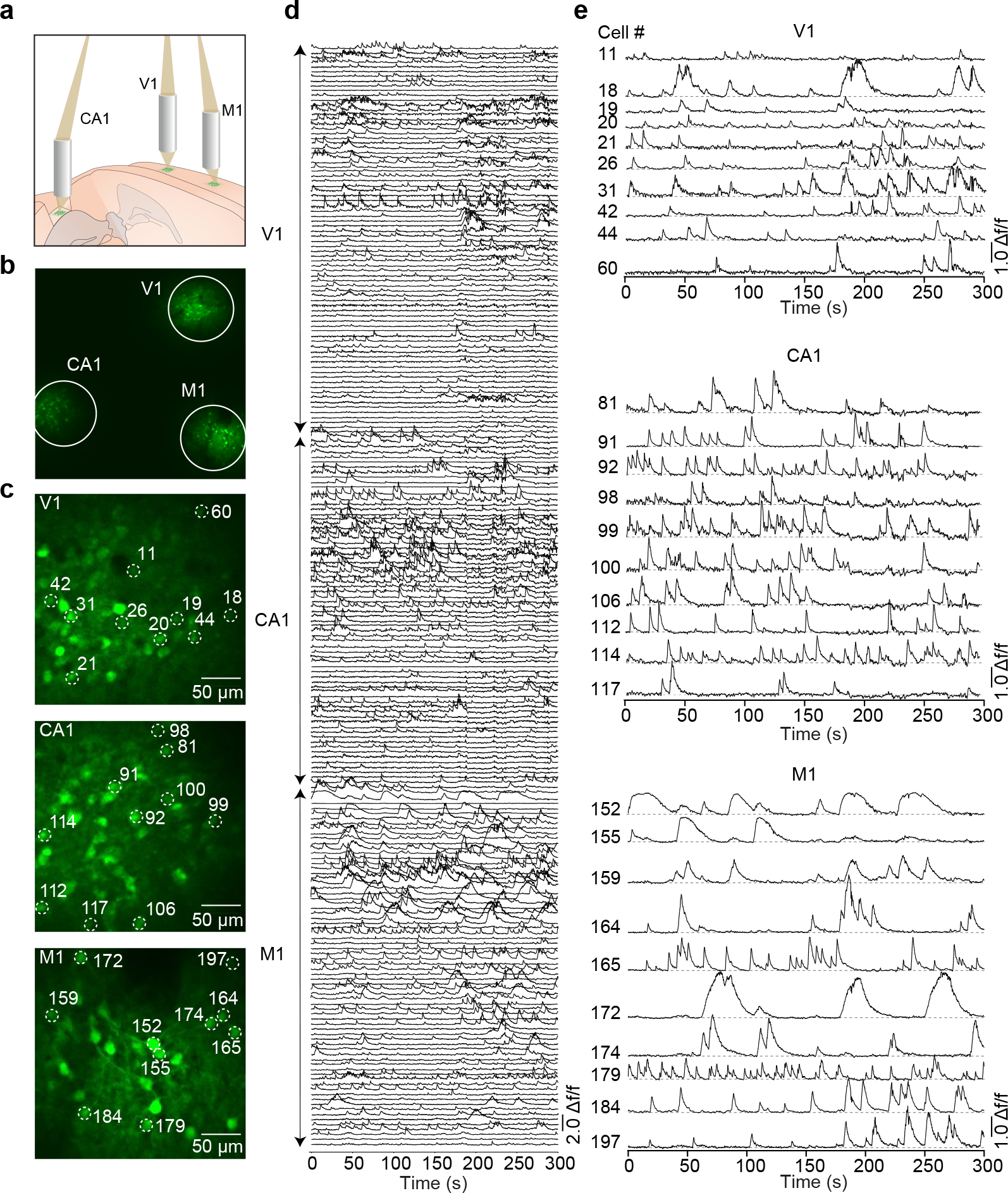
Application of MATRIEX imaging. **a**, Cartoon shows the configuration of MOs in V1, M1 and CA1. Note that the MO for hippocampus was different from the MOs for cortex, it was 0.25 mm shorter but has a 1.3 mm longer back-focal distance, thus compensated the depth difference between the cortex and the hippocampus. **b**, Full frame image of imaging areas in V1, M1 and CA1 (1200 × 1200 pixels per frame) of mouse, white solid circles outlining each area. **c**, Magnified images of each area. 10 neurons marked by white dashed circles in each imaging area were selected as examples to show later in panel e. **d**, Spontaneous activities (Ca^2+^ signal traces) of each neuron in each area imaged simultaneously for 300 s. Neurons were sorted by their relative locations. e, Enlarged view of the Ca^2+^ signal traces for the 10 example neurons marked in panel c.

In summary, we conclude that the MATRIEX method, based on the principle of multi-axis optical conjugation and two-stage magnification, has enabled two-photon Ca^2+^ imaging of neuronal population activities simultaneously in different brain regions at different depths (e.g., V1, M1, CA1) at single-cell resolution in awake mouse. There are many other alternative methods such as three-photon imaging^3, 17^ or fiber-based endoscopic imaging methods^18^ that can as well achieve deep-tissue in-vivo imaging. However, the uniqueness of the MATRIEX imaging method is the ability to simultaneously image in multiple brain areas that are at very different depths of more than 1 mm apart, which is so far almost unachievable by conventional microscope systems based on single optical axis.

A step-by-step device construction manual and a list of material cost is described in the Supplementary Methods with photo illustrations in Suppl. Fig. 3. Practical implementation of the MATRIEX method requires a very low material cost of less than $2,000 that is even less than many commercial microscope objectives for two-photon imaging. Importantly, it requires neither hardware modification nor software development on the existing conventional two-photon microscope system (in view that many two-photon microscopes belong to central facilities, can be easily accessed but cannot be easily modified), except for simply swapping the original microscope objective with the customized compound objective assembly. Furthermore, a broad range of other methods and techniques are compatible and may be further combined with the MATRIEX method to extend the applicability: (1) random-access scanning and/or stimulating methods^19–22^; (2) surgical techniques for chronic imaging^23^; (3) adaptive optics^24^; (4) head-mounted miniaturized scanning devices^25^. Thus, we expect that the application of the MATRIEX method will greatly advance the study of brain-wide neural circuit dynamics at single-cell resolution.

## Acknowledgements

The authors are grateful to Dr. I. Nelken, Dr. S. Remy and Dr. D. Kleinfeld for very helpful discussions; to Ms. J. Lou for help in composing the figures. This study was supported by the “100-Talents Program for Elite Engineers” of the CAS (H. Jia), the Key Scientific Research Equipment Development Project of the CAS (Super-resolution Microscopy Systems and Key Components, ZDYZ2013-1), the “1000-Talents Program for Young Scholars” of China (X. Chen), grants from the Ministry of Science and Technology of China (“973 Program”: 2015CB759500, 2018YFA0109600), the National Natural Science Foundation of China (61705251, 81671106, 81771175, 31700933, 81721001) and the China Postdoctoral Science Foundation (2018M632374), X. Chen is a junior fellow of the CAS Center for Excellence in Brain Science and Intelligence Technology.

## Author Contributions

X.C. and H.J. conceived the project. Z-Q.Z., M.Y., B.L., J.L, Y.G., Y.T., J-S.G., X.C. and H.J. designed and manufactured the optical instruments; M.Y., J.Z., T.L., K.Z, J.Y., Z-Q.Z., Z-M.Z., X.C. and H.J. designed and performed the biological experiments, M.Y., J.Z., T.L., J-H.G., X.L., Z.V., X.C. and H.J. analyzed the data with the help from all authors; X.C. and H.J. wrote the manuscript with the help from all authors.

## Author Declarations

All authors declare no competing financial interests and consent with the manuscript. All data in the manuscript are novel and have not been published elsewhere except that they will be uploaded to the preprint server bioRxiv. All technical details of the presented method are disclosed in online supplementary materials. Readers are welcome to comment on the online version of the paper and request technical materials or assistance for free.- Correspondence should be addressed to either of the following authors: Z-Q.Z., J-S.G. or X.C, technical materials and requests should be addressed to H.J.

