## Supplementary Figures and Methods for "MATRIEX Imaging: Multi-Area Two-photon Real-time In-vivo Explorer"

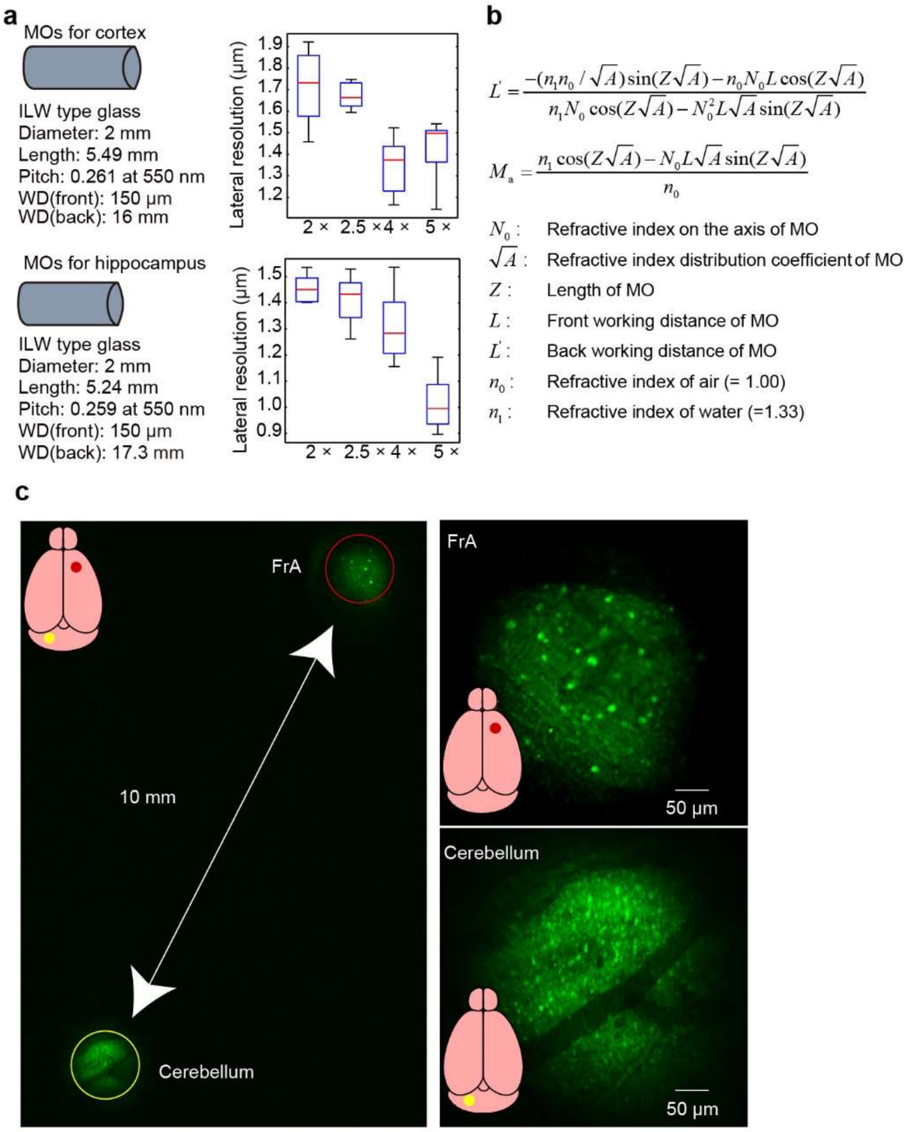


**Supplementary Figure 1 | Specifications of MOs. a,** Left icons and specifications: the two types of MOs designed and used in this study, for cortex and hippocampus, respectively. These parameters can be directly forwarded to the manufacturer to place orders. Graphs on the right show the lateral resolutions measured by using 0.51 μm beads for the two types of MOs in combination with four different models of AOs. **b**, Formula for calculating the angular magnification factor ($M_a$) of the MO. **c**, Two-photon image of the frontal association cortex (FrA) and the cerebellum of a GAD67-GFP mouse. Left image: full-frame scanned image by MATRIEX, upper-right and lower-right images: individually enlarged view of the two imaging areas. This experiment was to demonstrate the extreme case of far-away brain areas to be imaged by the MATRIEX method.

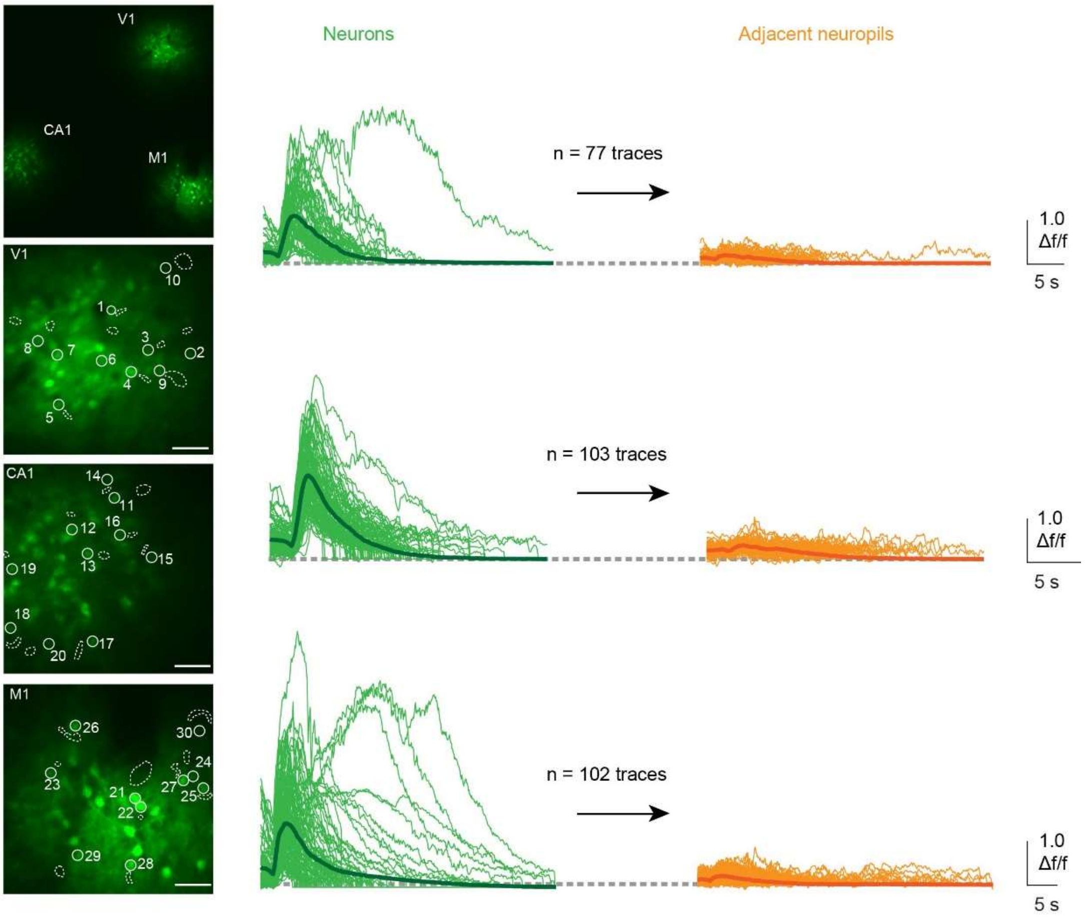


**Supplementary Figure 2 | Neuropil analysis.** For the sample experiment shown in Fig. 2 we extracted $Ca^{2+}$ signal traces from additional regions-of-interest (where there is no visible neuronal somata or obvious neuronal structure) that were adjacent to neurons in each imaging area and overlaid them to compare with the neuronal signal events (in the same time window). 10 example neurons were randomly chosen from each area and re-named (different from Fig.2). Scale bar: 50 μm

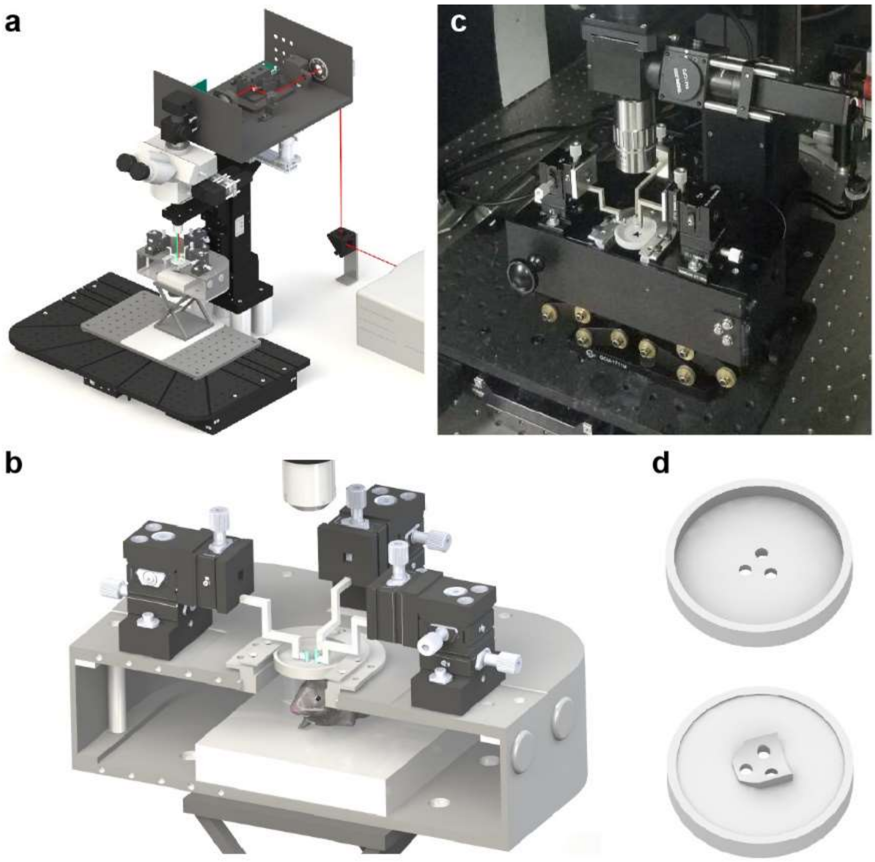


**Supplementary Figure 3 | Device design and construction. a,** CAD image of the entire MATRIEX imaging system including the conventional two-photon microscope. **b**, enlarged view of the compound objective assembly. **c**, Photo of the real device. **d**, CAD image of an example head chamber (2 views from top and from below). Note that the distance between the AO and the MOs is arbitrarily enlarged in order to see the micromanipulators.

**Supplementary Methods**

***MATRIEX imaging system.*** The MATRIEX imaging system consists of a standard upright two-photon microscope and a customized compound objective assembly. In this study we used a "LotosScan" microscope system built by the Suzhou Institute of Biomedical Engineering and Technology, Chinese Academy of Sciences, based on conventional single-beam resonant scanning technology same as our prototypes reported or used in earlier studies[1-7] as well as other commercially available products. A complete description of this microscope system is as below:

An ultrafast-laser (Fs Laser, Spectra-Physics MaiTai DeepSee) was used to deliver pulsed laser at ~100 fs pulse width, 80 MHz repetition rate at wavelength of 920 nm and average power of 2.3 W. The laser power fed into the scanner was modulated by a Pockels cell (PC, ConOptics model 1350-120 with driver M302RM). The laser beam was expanded by a beam expander (BE, Thorlabs GBE02-B) to fill the aperture of scanning unit. After the beam expander, the laser beam was raised to the scan box by a periscope. In the scan box, a mechanical shutter was used to stop the laser when the image acquisition was not running. The fast-axis scanner was a resonant mirror (SM1, Cambridge Technology, Model 12K CRS) and the slow-axis scanner was a galvanometric mirror (SM2, Cambridge Technology, Model 6215H). A custom-designed f-θ scan lens (SL) was used to flatten the scanning field of view in the imaging plane of objective lens. After the scan lens, a plane mirror was used to reflex the laser to the upright microscope body that has a tri-ocular port with built-in tube lens (TL, Olympus U-TR30IR). A dichroic mirror (DM, Semrock HC_735_LP) and a low-pass filter (Chroma ET700SP-2P8) were used to split the excitation and emission light. After passing a collector lens (Thorlabs AC254-30-A), fluorescence photons were detected by a GaAsP detector (PMT, Hamamatsu H10770PA-40) with over 40% quantum efficiency. The back pupil of the objective lens and the photosensitive surface of the PMT were conjugated by the collector lens such hat the loss of fluorescence was minimized.

A programmable data acquisition card (PXI-7851R, National Instruments) was used to control the parameters of scanner and PMT, and a high-speed data acquisition card (PXIe-5122, National Instruments) was used to digitize the PMT signal. Trigger bus of a PXI chassis (PXIe-1082, National Instruments) was used to synchronize the scanning trajectory and image acquisition. A customized LabVIEW software was developed to acquire and save live streaming image data. A reconstruction algorithm was used to correct the nonlinear scanning of the 12K resonance mirror. Time-lapse images can be taken either at 10 frames/s with 1200 × 1200 pixels or 20 frames/s with 600 × 600 pixels. The software features an online rolling average display function and several simple online analysis functions of $Ca^{2+}$ signal traces.. Original data was saved in a TDMS stream format and could be converted to TIF or AVI format by another customized LabVIEW program.

In the standard imaging mode, a standard water immersion objective (16x or 40x, Nikon) was used, same as in any conventional two-photon microscope system. In the MATRIEX imaging mode, the objective was replaced by the customized compound objective assembly. In principle, any upright two-photon microscope system that uses conventional infinity-corrected plan objectives, regardless of being custom-built or commercially purchased, can be directly used for implementing this method. There is no need to modify any hardware or software, except for replacing the microscope objective by the compound objective assembly. For each type of experiment, one must design and fabricate the miniature objective (MO), the head chambers and lens holders that together fit best. For the same type of experiment as shown in Fig. 2 we offer readers our custom-made head chamber and lens holders to test for free, whereas the MOs can be directly purchased from https://www.gofoton.com/ with the specifications as shown on Suppl. Fig. 1.

1. Locate the desired target areas on the brain atlas with stereotaxic coordinates.

   In principle, this method also works for large animals such as rats or marmosets, note that the accurate calculation of the coordinates is essential for the design of MOs because the different MOs need to be conjugated to the same virtual image plane. Tolerance of depth error is dependent on the AO, but typically less than 1 mm.

2. Get a proper air objective (AO)

   AOs with smaller magnification (e.g., 2x) can cover larger targeting zone, the diameter of which, can be estimated by an experience-derived formula: $D_{zone} = D_{scan}/M_{AO}$, where $D_{scan} \approx 24\ mm$ is the largest possible scanning field (depending on the two-photon microscope manufacturer). However, for imaging in mice, except for the extreme combination conditions (e.g., frontal cortex and cerebellum, Suppl. Fig. 1), AO of 4x magnification is sufficient to cover the targeting zone and with better resolution than the AO of 2x.

3. Work out an ideal angle for the virtual image plane and head chamber.

   In most cases, the virtual image plane can be simply set as a horizontal plane in the stereotaxic system at a certain height above the skull surface. This is also ideal solution for many cases because the head chamber will also be parallel to the animal skull surface thus the animal is not tilted. However, in case that some brain areas are very difficult to access from the vertical direction, it may be desirable to work out a geometric solution by tilting the virtual image plane at certain angle.

4. Design and fabricate MOs with different working distances for conjugating target areas at different depths.

Geometrically, any 3 points (center points of target image areas) that are not on one line can determine one plane in 3D space. In this view, users can choose to skip this step by working with uniform MOs but requiring more effort in designing head chamber and surgical procedures. Note that MOs can be recycled after each experiment, but careful cleaning and sterilization is necessary before next use.

5. Design and fabricate a head chamber that fits the skull curvature and the insertion sites of MOs.

   Based on the information of steps 3 and 4, design a head chamber for head-fixation of the animal under the microscope. 3D-printing technology is highly recommended. Note that each of the insertion channels on the head chamber must be a bit larger than the outer diameter of the MOs in order to have a certain room for mechanical manipulation. The head chamber is a consumable that is glued to the animal skull before surgery begin and disposed of together with the animal after experiment.

6. Test the assembly on a 3D-printed animal skull under surgical stereoscope

   We also recommend performing test experiments by mounting MOs with micromanipulators underneath a surgical stereoscope for an animal right after surgery, to directly visualize whether all MOs show best focused view of the target area (vasculatures, for example see Fig. 1d).

The above 6 steps can be iterated until best design to be achieved for each desired experiment. We recommend to start with uniform MOs for the first round and then to improve the design based on the test result, e.g., to make different MOs, or to choose a different virtual plane angle and insertion patch, etc.

We demonstrated acute imaging experiments here in this study. For chronic imaging experiments, we recommend gluing the MOs onto the head chamber once after the first imaging session is successful, and then the animal can be detached from the head fixation rig to go free, for subsequent imaging sessions only the AO need to be re-adjusted (to focus on the virtual image plane).

A list of required materials and their cost is given in the following table:

| Component | Cost ($) |
|---|---|
| Animal head fixation rig | 240 |
| Head chamber (consumable, 3D-printed, batch of 100 pieces) | 100 |
| MO holders | 30 X 3 = 90 |
| MOs | 70 X 3 = 210 |
| AO (Olympus 4X) | 140 |
| Manual micromanipulators (for 3 MOs) | 380 x 3 = 1140 |
| Total | 1920 |

***Animals.*** C57/BL6J adult male mice (8-10 weeks old) were used in this study. All animals were provided by the Laboratory Animal Center at the Third Military Medical University. All surgical tools and the MOs were sterile before use. All the experimental procedures were performed in accordance with institutional animal welfare guidelines, were approved by the Third Military Medical University Animal Care and Use Committee, and were similar to our earlier reports[8-11].

For the GCaMP6f labeling of neuronal populations in multiple areas, we performed multi-site virus injections. The C57/BL6J mouse was anesthetized with 1-1.5 % isoflurane in oxygen and placed in a stereotactic head frame on a heating pad (37.5–38 °C). After we injected 100 µL lidocaine (2%) and applied eyes with ointment (Bepanthen, Germany), we removed skin and cleaned skull by sterile cotton-tipped applicators, then several craniotomies were performed with a dental drill. We infused the virus (200 nL, AAV(2/8)-syn-GCAMP6f) through a glass micropipette with a tip diameter of 10-20 µm. The depth of imaging was about 150 µm beneath the cortical surface, therefore we inserted the pipette tip slowly at oblique angle in 60 degrees vertically at where the injection spots were -0.26 mm anteroposterior to the site of imaging. We injected virus vertically in the depth of 1.2 mm for CA1 imaging. The pipette was standing for 2 mins before retraction. The scalp incision was closed using tissue glue (3M Animal Care Products, Vetbond), and post-injection analgesics were provided for 3 days to help recovery. All the viruses were purchased from Shanghai Taitool Bioscience Corp Ltd China. The imaging experiments were performed during the time window of 8~10 weeks post-injection.

As for surgery before imaging experiments, the first few steps were the same as the virus injection procedure. we glued the customized chamber to the skull with UHU glue. Next, we performed craniotomies (slightly more than 2 mm diameter which is the outer diameter of the MO) at each area. It is strongly recommended that skull openings should be done after all craniotomies are completed, so to minimize the time of exposing brain tissue. Then, we performed aspiration to remove the tissue over the callosum and CA1. In case that bleeding happened, we used normal artificial cerebral-spinal fluid (ACSF) containing 125 mM NaCl, 4.5 mM KCl, 26 mM $NaHCO_3$, 1.25 mM $NaH_2PO_4$,

2 mM $CaCl_2$, 1 mM $MgCl_2$ and 20 mM glucose (pH=7.4 when bubbled with 95% oxygen and 5% $CO_2$) to swill blood out by droppers gently.

***Spatial resolution measurement***. We used 0.51 µm fluorescent beads (Bangs Laboratories, FC03F) for measuring axial and lateral resolutions and 20 µm fluorescent beads (Sicastar-greenF, 42-00-204) for calibrating the pixel sizes at different zooms. A 3D translation stage (Standa) holding the sample plate was used for measuring the travelling distances while image stacks are acquired. ImageJ was used for analyzing the image stacks to measure the full-width-at-half-maximum (FWHM) of the beads appearing on images and to assess the resolution.

***$Ca^{2+}$ imaging data analysis*.** We analyzed our data with using custom-written software in LabVIEW 2012 (National Instruments), Igor Pro 5.0 (Wavemetrics) and Matlab 2014a (Mathworks). To correct motion-associated artifacts in imaging data[12], we used a frame-by-frame alignment algorithm to minimize the sum of squared intensity differences between each frame image and a template which is the average of the selected image frames. To extract fluorescence signals, we visually identified neurons and performed the drawing of regions of interests (ROIs) based on fluorescence intensity. Fluorescence changes (f) were calculated by averaging the corresponding pixel values in each specified ROI. Relative fluorescence changes $\Delta f/f = (f-f_0)/f_0$ were calculated as $Ca^{2+}$ signals, where the baseline fluorescence $f_0$ was estimated as the 25th percentile of the entire fluorescence recording.
